# Effect of a constant magnetic field on morphology and motility of cell with cytoskeleton-associated magnetic nanoparticles

**DOI:** 10.1101/2024.05.12.593754

**Authors:** Olga Karavashkova, Alexandra Maltseva, Artem S Minin, Alexander Demin, Pavel Tin, Aleria Aitova, Valeria Tsvelaya, Anastasiia Alfredovna Latypova, Ilya Zubarev

**Affiliations:** Moscow Center for Advanced Studies; Institute of Metall Physics UrD RAS; Postovsky Institute of Organic Synthesis, Russian Academy of Sciences (Ural Branch)

## Abstract

Cell motility, shape supporting, and intracellular signaling are followed by changes in cell morphology and cytoskeleton. The cell reaction and the reorganization of the cytoskeleton occurs in a single volume of the cytoplasm and affects all components of the cytoskeleton: intermediate filaments, microtubules and microfilaments. A promising way to manipulate cells is magnetic nanoparticles that control cellular physiology. This approach is called magnetogenetics and has found application in various fields of cell and molecular biology. Using a magnetic field, it is possible to non-invasively regulate biochemical processes, migration and changes in the morphology of cells with magnetic nanoparticles. Our work opens up new possibilities for spatial manipulation of individual cytoskeletal components in vitro and operates biochemical pathways associated with individual cytoskeletal components.

## 2. Introduction

Magnetic nanoparticles are widely used in biological research and appear to be promising actuators of biological processes. Many articles propose magnetomechanical destruction of cancer cells using iron oxide nanoparticles and low-frequency rotating magnetic fields [10.1039/D1NA00474C]. At the same time, morphologically asymmetric rod-shaped MNPs can kill cancer cells more effectively than spherical MNPs when exposed to a magnetic field, due to their mechanical vibrations [10.1186/1556-276X-9-195]. Such works have not found application in clinical practice, but the use of magnetic nanoparticles can help in the study of many biological processes.

Magnetic nanoparticles used in the control of stem cell differentiation. Through mechanosensitive stimulation, it is possible to modulate osteogenesis [10.1007/s12274-016-1322-4 and chondrogenesis 10.3389/fbioe.2021.737132] in vitro. Interestingly, depending on the magnetic field of low or high intensity, the direction of differentiation (osteogenic or adipogenic) can be determined [10.1016/j.msec.2020.110652]. The influence of magnetic nanoparticles on cell fate and on the control of stem cell differentiation can be realized through mechanosensitive signaling pathways [10.1007/s10439-017-1884-7]. Mechanical stress from magnetic nanoparticles results in cell tension and partly reproduces in vivo conditions [10.1016/j.bioadv.2022.213028]. Thus, it is even proposed to mechanically influence the regulation of the function of neuronal cells [10.3389/fnins.2018.00299 and even stimulate the differentiation of stem cells in the neuronal direction 10.1007/s12015-022-10332-0]. Magnetic nanoparticle-driven activation drives asymmetric tissue growth and proliferation, leading to enhanced patterning in human nerve cells [10.1038/s41467-023-41037-8].

Using magnetic nanoparticles and electromagnetic tweezers, it is possible in vitro to determine the viscosity of the cytoplasm of a living cell, which is important for modeling the behavior of nanoparticles in cells and determining the physical characteristics of cells [10.3390/ph16020200].

The action of a magnetic field and magnetic nanoparticles can non-contactly influence cell motility and the direction of migration [10.2147/NSA.S298003]. In this way, we can set the location of cells on the plastic, regulate the speed and direction of migration [10.1039/C6QM00219F] due to the traction force on the periphery of the cell [10.3390/ijms21186560]. This directional cell migration is particularly important for the directional growth of neurons and Schwann cells [10.2147/IJN.S227328]. Some papers have postulated the use of magnetic nanoparticles to generate and spatially manipulate magnetic multicellular spheroids [10.1016/j.biomaterials.2009.12.047].

In studying the physical and biochemical properties of cells, it is important to obtain information about the response of individual cell components to physical influence. Magnetic nanoparticles can nonspecifically bind and remodel the cytoskeleton, which contributes to changes in the mechanical properties (rigidity) of cells [10.1186/s12951-021-00790-y]. The association of nanoparticles with the cytoskeleton can be used to selectively treat cancer cells by disrupting the cytoskeleton [10.1038/srep33560] or whole cells and cell-cell contacts [10.3390/ijms21249350]. Many methods and materials have been proposed to disrupt the cytoskeleton with various nanoparticles [10.1016/j.bioactmat.2023.06.016]. By encapsulating magnetic cobalt-platinum nanoparticles inside microtubules, magnetically induced alignment of microtubules outside cells was achieved [10.1021/acs.nanolett.0c01573]. At the same time, magnetic nanoparticles have not been used to study fundamental problems in the biology of the cytoskeleton. In our work, we are the first to propose an approach to selective manipulation of various components of the cytoskeleton (intermediate filaments, microfilaments, microtubules) to study cell behavior and clarify the role of individual components in the processes of spreading and migration.

## 3 Materials and Methods

### Nanoparticles

The article used magnetic nanoparticles of the composition iron core - carbon shell (Fe@C) and Fe3O4 nanoparticles, modified with functional groups that provide hydrophilicity and cross-linking with proteins.

Iron-carbon magnetic nanoparticles were synthesized and modified in the Laboratory of Applied Magnetism, Ural Branch of the Russian Academy of Sciences, Institute of Physics and Metallurgy using the gas-phase technique [https://doi.org/10.1016/j.carbon.2014.03.034]. In this case, the iron wire was evaporated in a flow of inert gas (argon) with an admixture of hydrocarbons (butane, propane, isobutane) by a high-frequency alternating electromagnetic field. On molten iron nanodroplets, catalytic decomposition of hydrocarbons occurred with the formation of a carbon shell, after which the nanoparticles were captured by a fabric filter. Surface modification using aryl-diazonium derivatives [10.1134/S1995078010070037], [https://doi.org/10.1016/j.colsurfb.2019.01.009], for this, nanoparticles were mixed with the diazonium derivative 4-aminophenylacetic acid (Sigma Aldrich, USA) and were processed for 30 minutes using a submersible ultrasonic activator, after which they were separated using a magnet and washed. The saturation magnetization is 90 emu/g, the size according to TEM data is ∼10 nm.

MNPs based on Fe3O4 were synthesized by analogy with the articles [https://doi.org/10.1016/j.ceramint.2021.04.310], [https://doi.org/10.1021/acsami.1c07748]. Briefly, a saturated aqueous solution of ammonia was added to a solution of FeSO4 × 7H2O and FeCl3 × 6H2O in H2O with stirring and sonication in an ultrasonic bath (40 °C). After 10 min, the particles were deposited on a magnet and washed with H2O to neutral pH and dispersed in H2O. To functionalize the surface, a solution of PMIDA in H2O was added to the MNPs. The colloidal solution was mixed using an overhead stirrer. The precipitation of functionalized MNPs was carried out in a centrifuge at 25 thousand rpm for 15 min; the particles were washed with H2O and dispersed in 50 ml of H2O.

The nanoparticle suspension was filtered through 0.22 μm PTFE syringe filters (JetBiofill, China) and diluted with nutrient medium to the required concentration.

### Carbodiimide crosslinking nanoparticles with antibody

To target delivery of carboxylated nanoparticles to cytoskeletal proteins, we modified the surface of NPs with antibodies using the carbodiimide conjugation method [https://doi.org/10.1016/j.jconrel.2020.01.035]. Primary antibodies to vimentin (Cell Marque, 347R-25), polyclonal antibodies to beta-actin (ThermoFisher, PA1-183) and antibodies to acetylated tubulin (Sigma-Aldrich, T7451) were used. Ig G+H with a fluorophore AF-488 (ThermoFisher, A-11013) was used as secondary antibodies.

To activate carboxyl groups on nanoparticles, EDC (Sigma-Aldrich, 39391) and NHS (Sigma-Aldrich, 130672) were used in a mass ratio of 1:3 and a mass ratio of EDC to nanoparticles of 6:23. The activation reaction of nanoparticles pretreated with an ultrasonic dispersant was carried out at room temperature for 20 minutes in MES buffer (pH 6,0). To remove reaction byproducts and free EDC and NHS molecules, the mixture was centrifuged for 5 minutes at 21,130 rcf at room temperature, after which the supernatant was removed. Buffer and antibodies were added to the sediment in ratios of 1:23 - 2:23 to the volume of nanoparticles and incubated for 60 minutes at room temperature. The carboxyl groups of the nanoparticles formed an amide bond with the primary amino group of the antibodies. To remove unbound components from the reaction solution, the mixture was centrifuged for 5 minutes at 21130 rcf, the supernatant was removed, and the modified nanoparticles were diluted with serum-free culture medium.

### Crosslinking of nanoparticles with protein G/L and Cy5 fluorophores

To 1 mg of nanoparticles, 1 mg of EDC and 1 mg of NHS were added in 100 μl of 0.1 M MES, pH 6.0. The solution was immersed in an ultrasonic bath and incubated for 30 minutes at RT. 1 mg of protein G/L in PBS and 100 μl of borate buffer pH 8.0 were added, the tube with the reaction mixture was placed in an ultrasonic bath and incubated for 30 min at RT. Next, Cy5 fluorophore 20 μg was added, and the solution was left for 8 hours or overnight. To remove contaminants from the solution the mixture was centrifuged for 10\ minutes at 21,130 rcf at room temperature, after which the supernatant was removed. This process was then repeated twice.

### Paper chromatography of nanoparticles with antibodies

The volumes of nanoparticles used were varied: 2.5 μl and 6.25 μl, antibodies were used: 1, 4 and 10 μl, and the crosslinking time was varied (30, 60 minutes).

Antigen (BSA, Sigma-Aldrich, A8022) was applied to a membrane made of nitrocellulose and an absorbent pad (UniSart CN 140) [https://doi.org/10.1016/j.jcis.2019.01.065] in the form of a drop of ∼2.5 μl 10 % solution (Figure in supplement). The mobile phase was prepared from 8 μl of a solution of nanoparticles cross-linked with antibodies to BSA (ThermoFisher, A11133), 4 μl of casein solution (0.2 mg/ml, Sigma-Aldrich C3400) and 8 μl of PBS pH 7.4 (Sigma-Aldrich, P4474). The membrane was lowered into the mobile phase, which moved up along it within 30 seconds and reached the area with BSA, where BSA Antibody reacted with the antigen.

Due to the dark color of the nanoparticles, it is possible to visually detect bands on a chromatographic strip. Thus, the thickness of the band will correlate with the number of reacted NP-antibody complexes with the antigen and, accordingly, the preservation of the affinity of the antibodies.

### Staining of the cytoskeleton with the NP/antibody/dye complex

To visualize cytoskeletal components, AF-430 dye and beta-actin antibodies cross-linked to iron nanoparticles via protein L (ArtBioTech, 120423) were used. Two experimental options were tested: labeling with a complex with antibodies or separately. In the first case nanoparticles with protein L and AF-430 dye are cross-linked with antibodies, and then cells are incubated with an obtained complex. The second option suggests that cells are labeled sequentially with primary antibodies and then with nanoparticles with protein L and AF-430 dye.

To produce complexes, 0.3 μg of antibodies per 50 μg n/h was added to a solution of nanoparticles with protein and dye and incubated for 30 min at RT without further purification. Incubation of cells with complexes (nanoparticle/antibody/dye), as well as with pure antibodies, lasted 1 hour at 37 C, after which the cells were washed with PBS: 200 μl of PBS was applied to the well of a 48-well plate for 2 minutes, after which it was removed, and the operation repeated twice more. Finally, NPs/protein L/dye (labeled separately) were added to the wells containing only pure a/t, incubated for 15 min RT and washed three times with PBS.

At the 2nd stage, labeling was carried out in confocal dishes with a working volume of 1 ml. Primary antibodies to beta-actin, vimentin and tubulin were used.

### Magnetic systems

The article used magnetic systems with a vertical gradient, which make it possible to enhance the process of absorption of magnetic nanoparticles by cells, as well as magnetic systems with a lateral gradient, causing a force to accelerate the migration of cells with magnetic nanoparticles.

The vertical gradient magnetic system was made of cylindrical permanent NdFeB magnets with a diameter of 50 mm and a height of 20 mm. There is a plastic protective cover (1) on top that allows you to center the Petri dish on the magnet, which is bolted to the plastic body (2). Inside there is a cylindrical magnet (3) and under it a magnetic field concentrator of similar size (4) made of a soft magnetic alloy, which makes it possible to make the field more uniform along the edges of the magnetic system.

**Figure 1.**
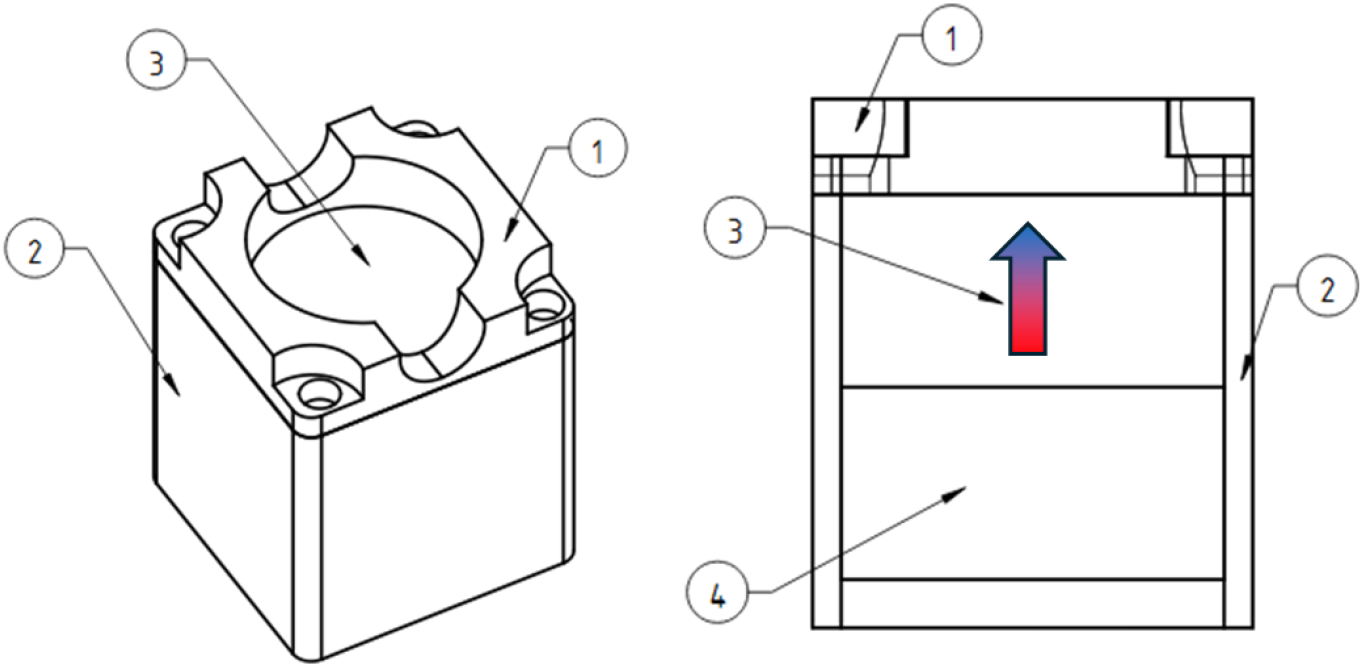
Diagram of the design of magnetic systems with a vertical gradient, the arrow indicates the direction of magnetization of the magnets. Explanations in the text.

The 3D printed plastic case does not have any structural load and is necessary for the safety of users (Figure A). Magnets of this size are attracted to each other and to ferromagnetic materials with great force and can seriously injure a person if used carelessly. The measured topology of the magnetic field is shown in (Figure B), the vertical field gradient is about 1E4 T/m.

**Figure 2.**
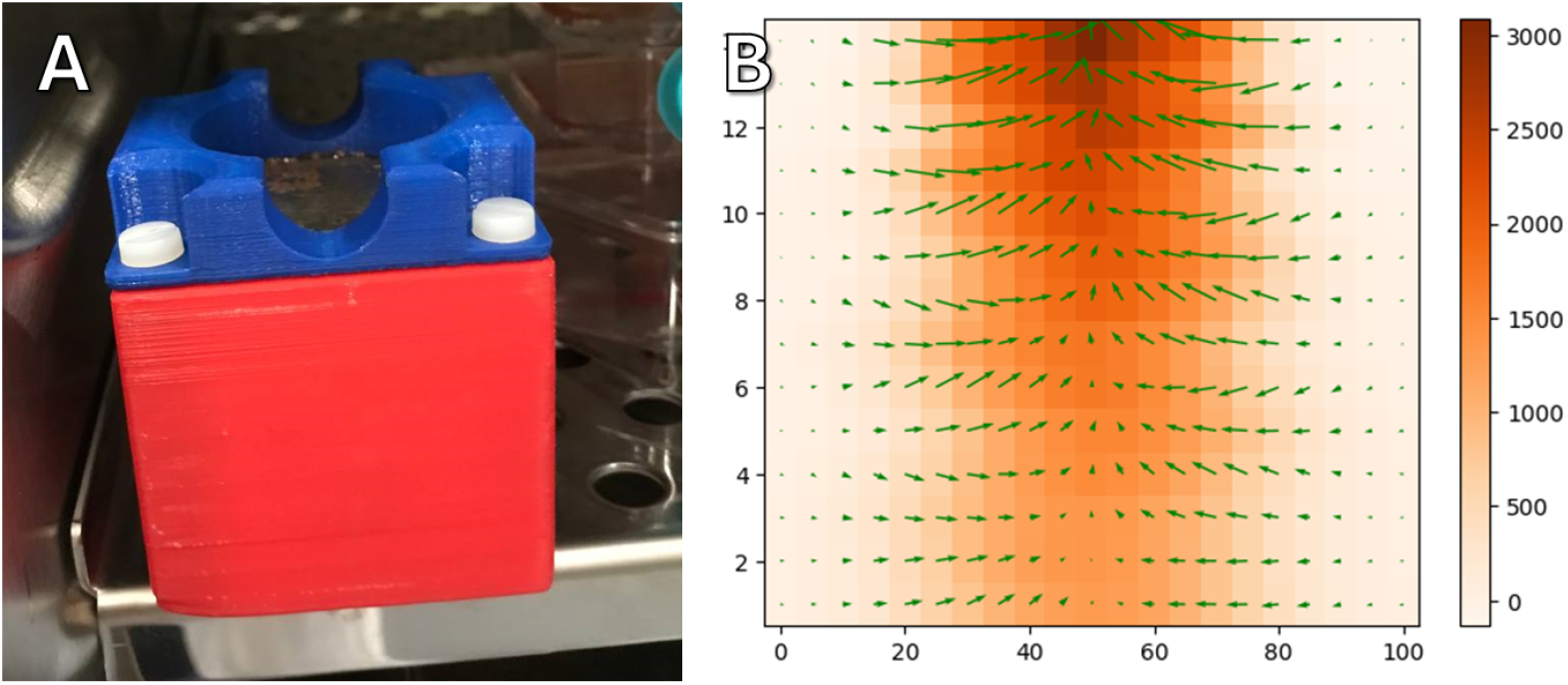
Photograph of the assembled magnetic system (A) in a cell incubator and measured topography of the magnetic field in the work area (values are given in Oe), green arrows indicate the direction of the field gradient. (B)

Lateral gradient magnetic systems were made from beveled shape permanent NdFeB magnets. This shape is designed to create a magnetic field with a uniform gradient to achieve uniform exposure to cells in culture. A sketch of the magnetic system is shown in the figure below. A petri dish (1) with cells is placed on a plastic holder (2) located between two magnets (3). Above and below the magnets there are magnetic field concentrators made of soft magnetic iron alloy (4). The oppositely directed magnetic field leads to the appearance of a repulsive force between the magnets, so that the magnetic system maintains its integrity; it is held externally by an aluminum housing (5).

**Figure 3.**
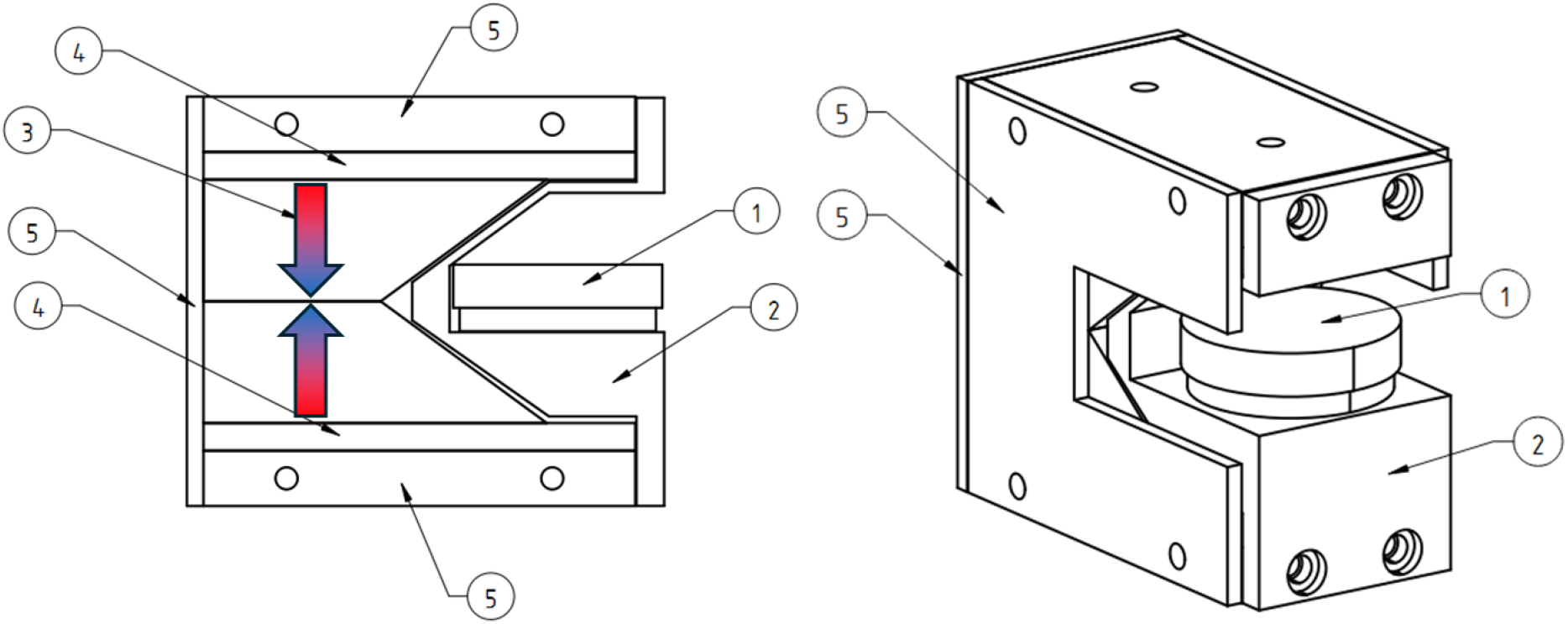
Diagram of the design of magnetic systems with a lateral gradient, the arrow indicates the direction of magnetization of the magnets. Explanations in the text.

The magnetic system is held together by screws screwed into the metal parts of the structure (Figure 4A). Using a Hall sensor, the topography of the magnetic field was taken in the working area of the magnetic system at the height where the dish with cells is located (Figure 4B). The measured gradient was 3*E4 T/m.

**Figure 4.**
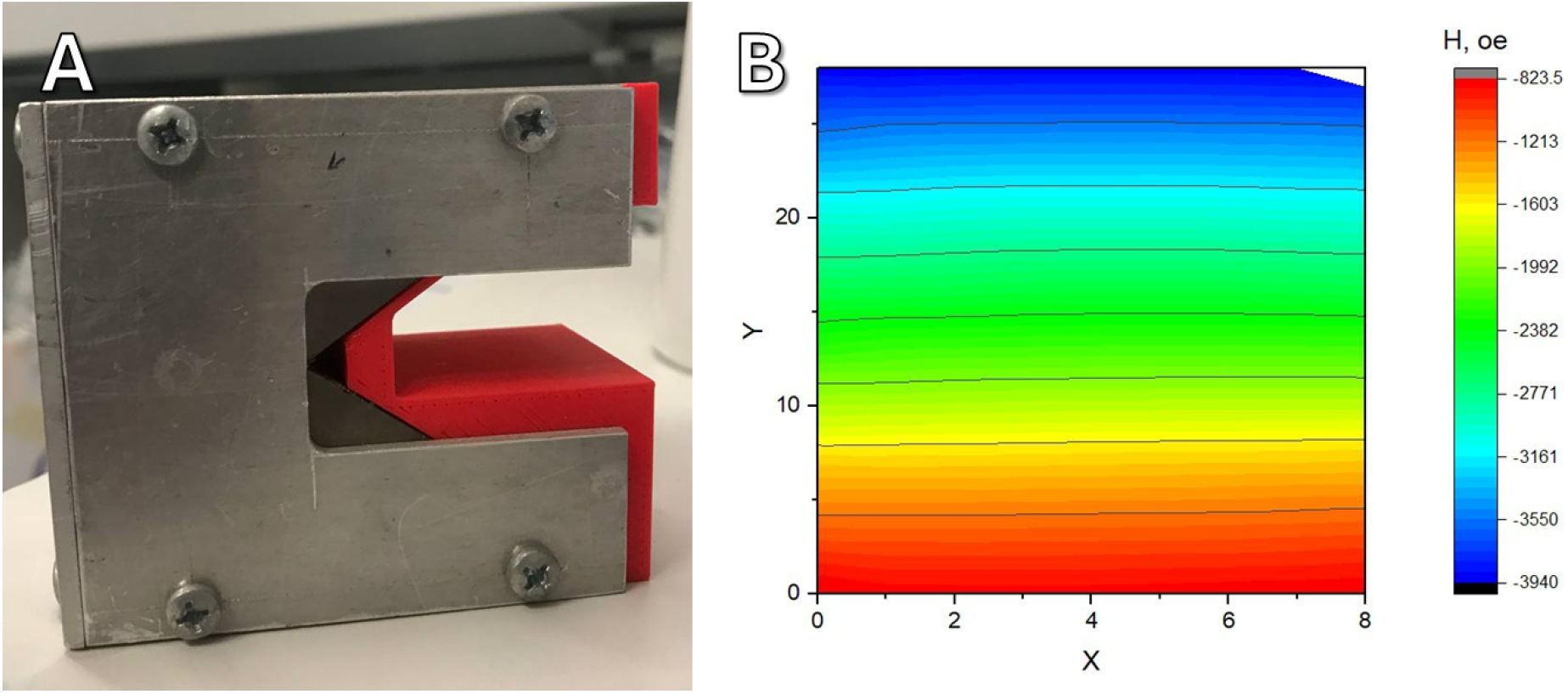
Photograph of the assembled magnetic system (A) and measured topography of the magnetic field in the work area (B)

### Cell culture

Experiments were performed on hTert MSC ASC52telo cell line obtained from Cell Bank of MSU (Russia). Cells were maintained on a commercially available cultural media DMEM (Paneco, P410п) with 10% FBS (Biosera), L-Glutamine, 100 U/ml penicillin, and 100 μg/ml streptomycin (Gibco). Cells were cultured in T25 flasks (Jet Biofil, China) or confocal Petry dishes (Jet Biofil, China) in an incubator at a temperature of 37°C in a humidified atmosphere with 5% CO2. Experiments were performed on cells with confluence levels from 70% to 100%. Trypsin-EDTA 0.25% solution (Paneco, п036п) and Versene solution (Paneco, P080п) were used for cell subculturing.

### Agarose molds

To study migration, cells were cultured in special agarose molds. Agarose moulds were made according to the method partially described in [https://doi.org/10.3390/ijms24098150]. For this purpose, master-forms were printed using MSLA 3d printer Photon Mono (Anycubic, China), after which they were filled with two-component cast silicone. After the silicone solidified, the forms were washed with isopropyl alcohol and further stored in zip-lock bags.

To make agarose molds, a molten solution of 2% agarose in PBS was poured into the silicone mold. After solidification (5-10 minutes depending on ambient temperature), the agarose mold (Figure) was transferred to a dish/well of a 6 well plate. The agarose mold was soaked in full nutrient medium for 15 minutes, after which it was ready for use (Figure).

**Figure 5.**
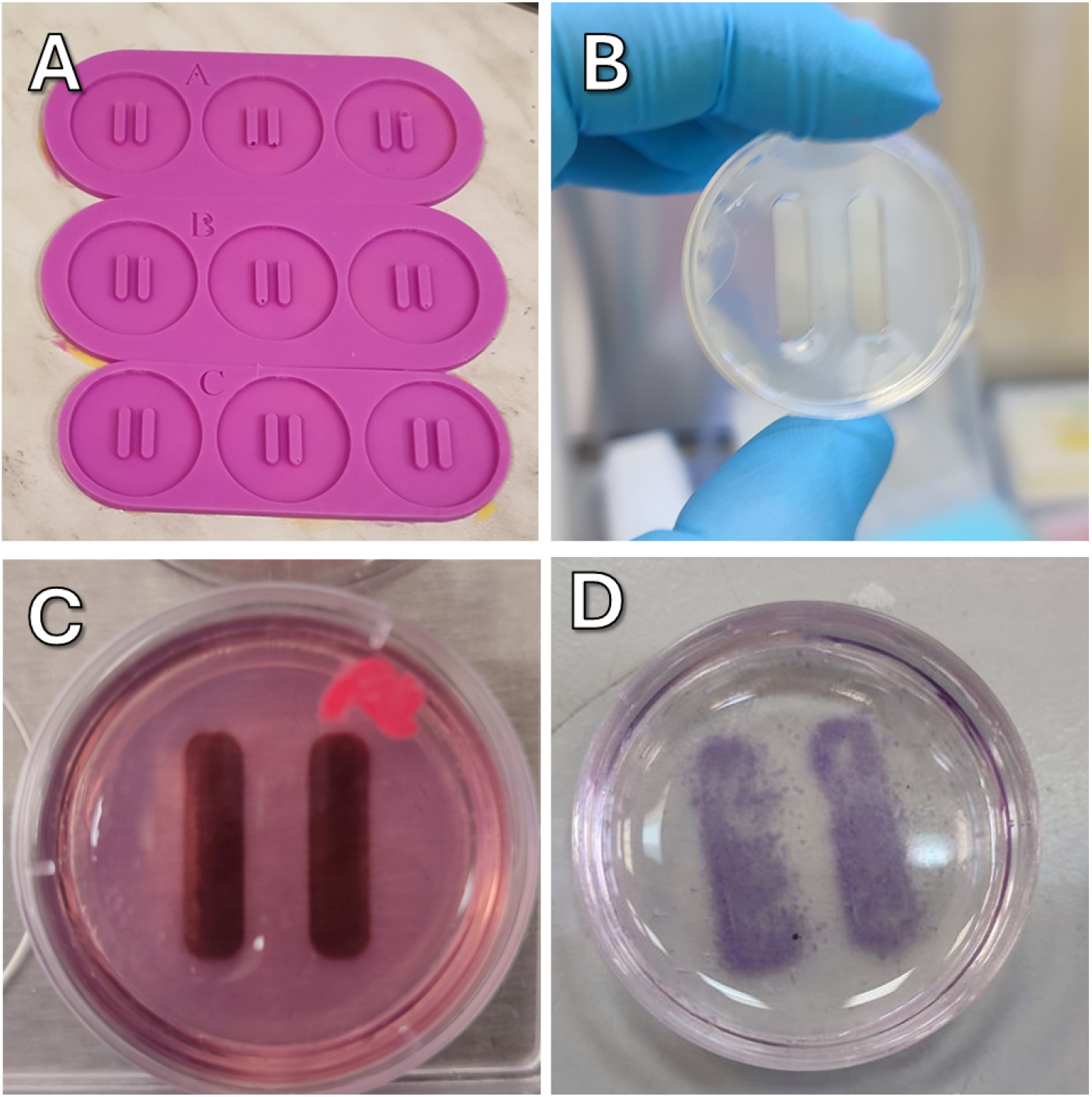
A - silicone molds, B - agarose mold mounted in a petri dish, C - suspension of nanoparticle-loaded cells poured into the cells of agarose mold, D - cells after extraction of the mold (H&E staining)

### Measurement of nanoparticle concentration in cells using NMR relaxometry

The process of uptake of nanoparticles by cells was studied using NMR relaxometry, using the technique described in [https://doi.org/10.1134/S0031918X19130027 and 10.1088/1742-6596/1389/1/012069]. NMR relaxometry with hydrogen protons allows one to measure the relaxation time of water, which changes significantly in the presence of ferromagnetic objects. Cells occupy an extremely small part of the liquid volume, so it is not possible to directly measure the number of particles captured by them. Therefore, measuring the residual concentration in the nutrient medium makes it possible to reliably calculate exactly how many particles have entered the cells. Quantitative measurements are possible provided that the nanoparticles are relatively stable in the environment and do not agglomerate. This affordable and effective method allows you to measure particle concentrations with sufficient sensitivity.

Cells were seeded at 70% confluency into a culture vessel and incubated for 24 hours. Next, the medium was changed to fresh medium containing nanoparticles; after a certain time, samples were taken (100 μl each) and their relaxation time was measured using an NMR relaxometer. To ensure a uniform distribution of nanoparticles that could settle over time, each time a sample was taken, the medium was intensively aspirated with a dispenser. The value of 1/T2 is linearly proportional to the concentration of magnetic nanoparticles in the liquid, therefore this value is shown in the graphs [https://doi.org/10.1016/j.colsurfb.2019.01.009].

### Lysis protocol

After removing the cells from the plastic using trypsin, the cell suspension tubes were placed in an ultrasonic bath for 30 seconds (Fisherbrand, FB11203) to partially disrupt the cell membranes and make the cytoskeletal fibrils more accessible. Antibodies to various cytoskeletal proteins (vimentin, actin and tubulin), cross-linked with iron oxide nanoparticles. (5 μl of NPs solution (2,3 mg/ml) was mixed with to 2 μl of antibodies stock solution in 50 μl PBS) were added to the resulting suspension; the mixture was incubated in a thermostat at 37°C for 30 minutes, after which secondary antibodies labeled with a fluorophore AF-488 were added. The resulting suspension was carefully washed using magnetic separation. The test tube was placed on a magnetic separating stand, cell fragments associated with nanoparticles were magnetized to the side for 1-2 minutes, the remaining liquid was carefully removed with a dispenser and replaced with 100 μl of PBS and placed in a magnetic system with a lateral magnetic field.

### Cells with magnetic nanoparticles in vertical gradient magnetic and lateral gradient magnetic systems

To accelerate the uptake of nanoparticles by cells and bypass lysosomal uptake, vertical gradient magnetic systems were used. A solution of nanoparticles in the required concentration, with the modification with any protein, in a culture medium without FBS was added to a Petri dish with cells. To suppress vesicular transport, the endocytosis inhibitor dansylcadaverine (Sigma-Aldrich, D4008) was used. The cells were incubated with the nanoparticle solution for one hour, after which the solution was collected with an aspirator, the cell layer was gently washed with complete culture medium, and the medium was replaced with complete medium with 10% FBS. After half an hour in the incubator, the Petri dish was placed on a lateral gradient magnetic system with a marker indicating the orientation of the dish. On a side magnetic system, dishes with cells were incubated for 20 hours. Antibodies to Integrin beta-1 (Invitrogen, MA5-27900) were used to suppress cell migration.

## Results

### Analysis of the preservation of antibody affinity after cross-linking with nanoparticles

To assess the integrity of antibody affinity, we conducted a series of experiments. Using paper chromatography, we have shown that different ratios of antibodies to BSA and iron nanoparticles pass differently in the chromatographic strip depending on the ratio of the components and their quantity. The presence of bends at the site of application of the BSA protein indicates specific binding to the complex of nanoparticles and antibodies to BSA. No bands were detected on control membranes. Due to the formation of aggregates of iron nanoparticles in solution, only a small portion of antibody-bound nanoparticles move in the mobile phase, which may indicate low efficiency of direct cross-linking of antibodies and nanoparticles. When directly cross-linking antibodies with nanoparticles, it was shown that an increase in the amount of antibodies reduces the intensity of the bend, which can be explained by intermolecular cross-linking of proteins on the surface of nanoparticles when there is an excess of antibodies. When incubated for half an hour, cross-linking 6.25 μl of n/h with 10 μl of a/t turned out to be more effective than cross-linking with 4 μl, however, when the duration was increased to 1 hour, the intensity of the bands became equal (Figure in supplement). This indicates that less a/t was bound in half an hour than could have occurred at a given a/t concentration (4 μl), and increasing the crosslinking time allowed more a/t to specifically bind to the antibodies. Subsequently, due to the high cost of primary a/t, cross-linking was carried out with a smaller amount of a/t (4 μl) and within 1 hour without loss of efficiency and an amount of a/t of 6.25 μl.

Using this method, we showed that antibodies retain their binding affinity to their antigens after chemical cross-linking. Using direct cross-linking of antibodies with nanoparticles and dye, we were unable to obtain images of cells suitable for microscopy. In this regard, an indirect crosslinking method through additional antibody recognition proteins was proposed.

To detect cytoskeletal components in vitro, we proposed cross-linking nanoparticles with protein G or L. These proteins are able to recognize and strongly bind antibody chains [10.3390/ma9120994]. Nanoparticles were cross-linked with dye and protein G/L, and then conjugated with any antibodies to cytoskeletal components. In an experiment with antibodies to actin, it was shown that the resulting complex is able to bind and detect actin microfilaments. Labeling separately did not show good results.

**Figure 6.**
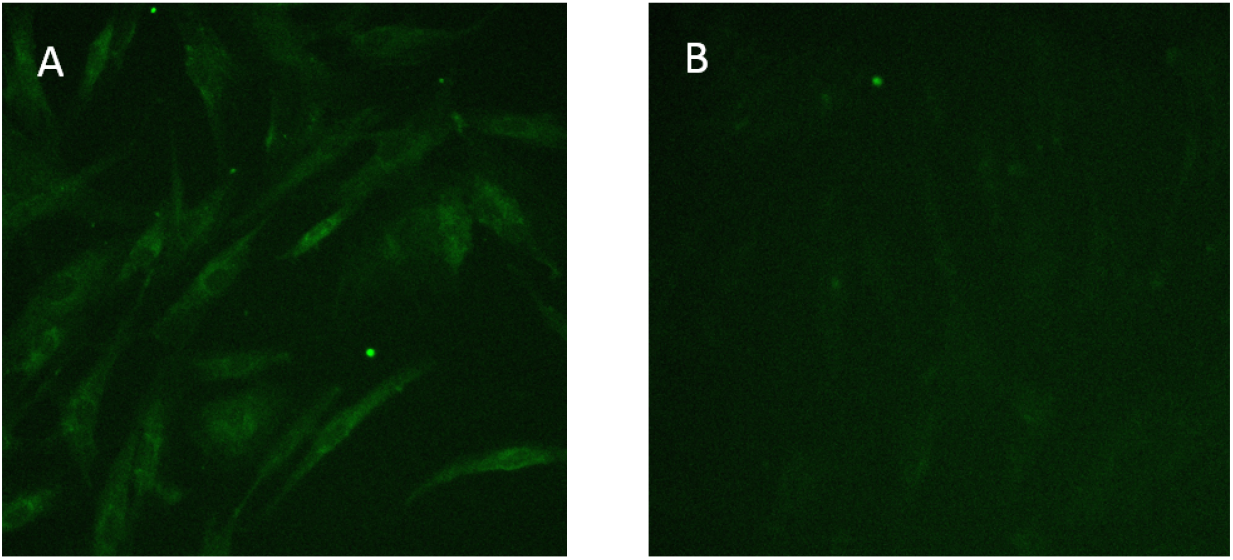
Immunocytolabeling with antibodies to actin. A: labeling with complexes with antibodies; B: labeling separately.

Similarly, it has been shown that vimentin filaments and microtubules can be associated and detected (Figure 7)

**Figure 7.**
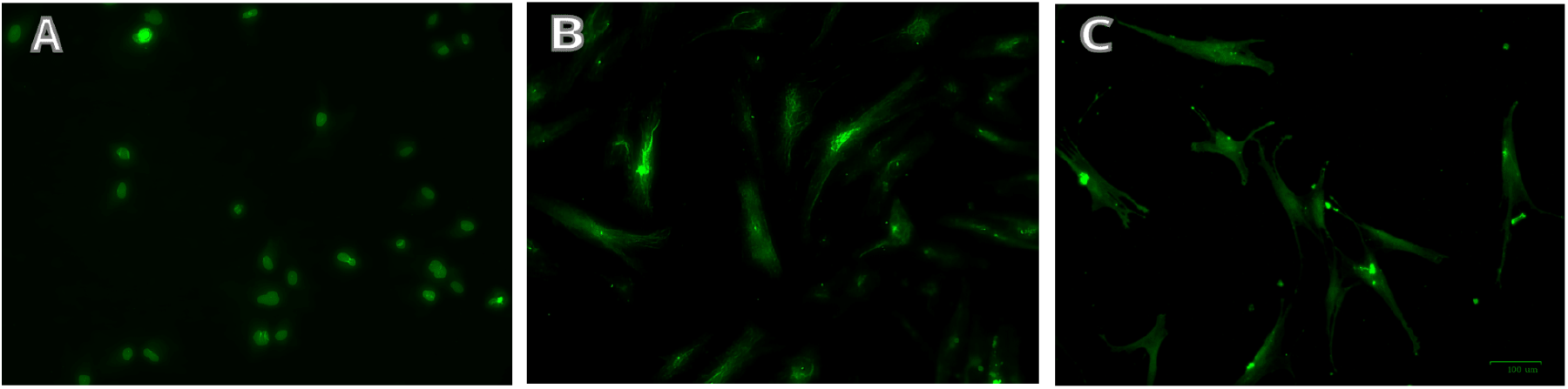
Immunocytolabeling. A: antibodies to vimentin; B: antibodies to alpha-acetyl-tubulin, C: antibodies to beta-actin.

### Using a vertical gradient magnetic field for the uptake of nanoparticles by cells

The uptake of magnetic nanoparticles by cells is not a rapid process, and with active uptake, a considerable part of the nanoparticles can accumulate in lysosomes, which prevents their binding to other cell components [https://doi.org/10.1039/C6CS00636A]. Therefore, accelerating the uptake of nanoparticles can reduce the time spent on experiments and avoid their accumulation in lysosomes. In order to accelerate the uptake of magnetic nanoparticles by cells, we applied a magnetic field with a vertical gradient, “dragging” the nanoparticles inside the cells. This process was monitored using NMR relaxometry. The figure below shows the change in the relaxation time T2 of hydrogen protons in the nutrient medium, presented as 1/T2, which is linearly related to the concentration of nanoparticles in the liquid. Blank experiments without cells were also carried out and the absorption of nanoparticles on culture plastic does not make a noticeable contribution to this process; all changes in the concentration of nanoparticles are associated with their active uptake by cells. When magnetic nanoparticles are added to the nutrient medium in high concentrations (on the order of 0.1-0.5 mg/ml in cells, they are visually visible (Figures 8B and 8C), however, in small concentrations, only NMR relaxometry allows monitoring the uptake of nanoparticles.

**Figure 8.**
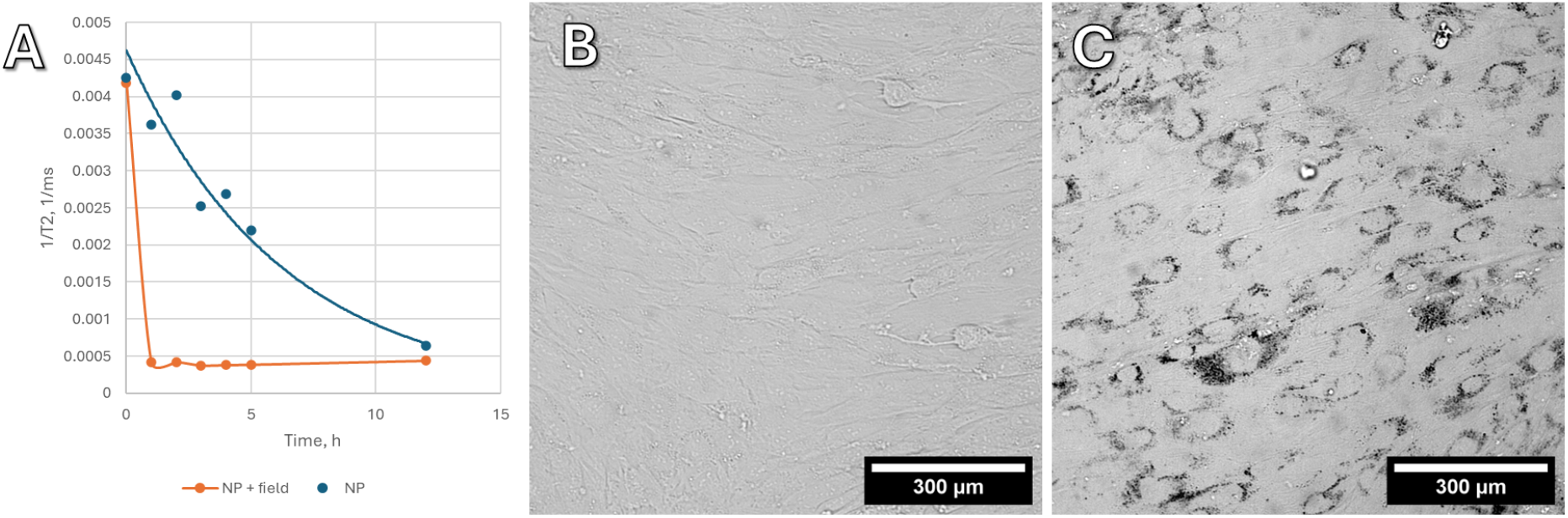
The process of uptake of magnetic Fe3O4 nanoparticles modified with carboxyl groups by hTERT culture cells.

### Lysis experiment

To confirm that after cross-linking with nanoparticles, antibodies retain the ability to bind to the corresponding antigens inside a living cell, experiments were carried out with lysed cells. Unlike a whole cell, in which the viscosity of the cytoplasm, the mechanical properties of the plasmalemma, and the spatial organization can prevent binding to the cytoskeleton and movement towards the magnet, in the cell lysate specific binding and pulling towards the magnet can be clearly demonstrated. We have shown that in cell lysate, antibodies cross-linked with nanoparticles retain their affinity and are specifically associated with the cytoskeleton. In the presence of a magnetic field, magnetization of fluorescently labeled antibodies bound to magnetic nanoparticles to the edge of the droplet near the magnet is observed (Figure 9). Interestingly, probably due to the greater stability of intermediate filaments with the same number of cells, the abundance of fluorescent aggregates in the experiment with vimentin is greater than in the experiment with actin and tubulin (Figure 9A).

**Figure 9.**
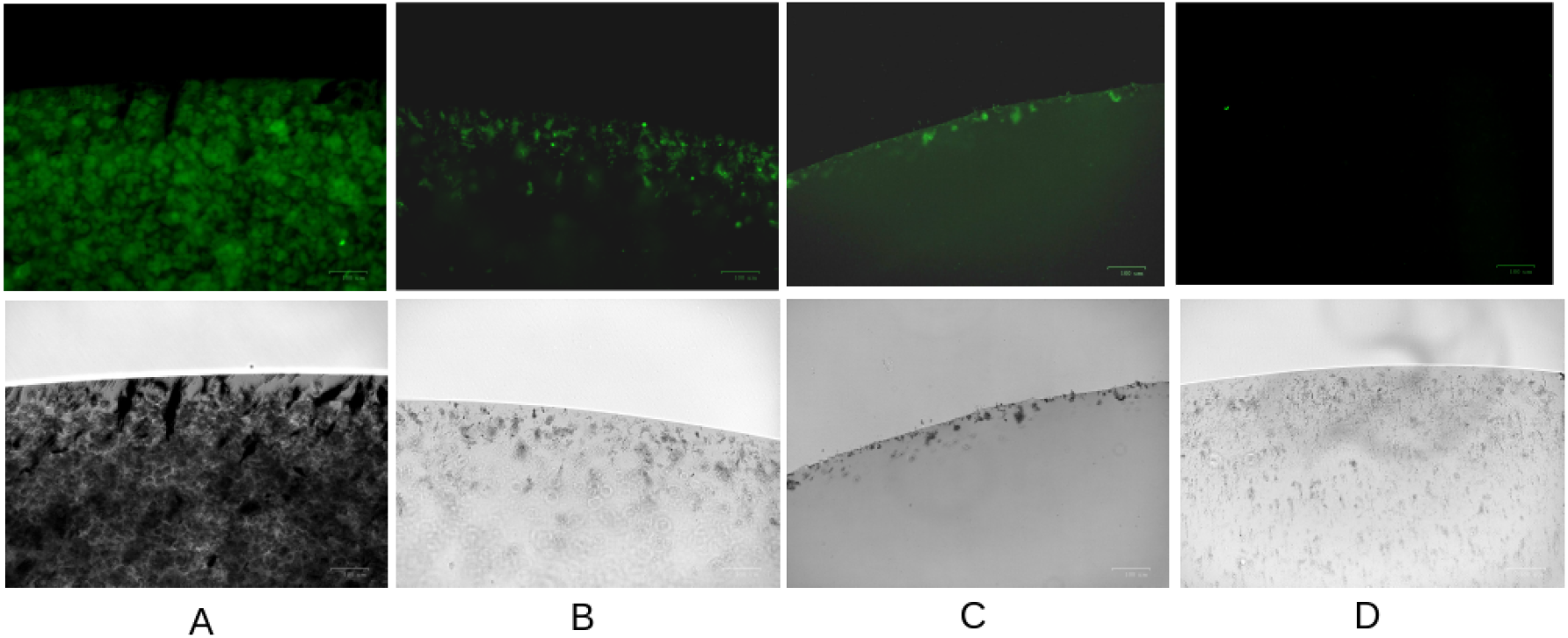
Lysis experiment. A antibodies to alpha-acetyl-tubulin, B - antibodies to vimentin, C - antibodies to actin, D - control without antibodies.

In experiments on the uptake of Fe@C and Fe3O4 nanoparticles into cells, and the experiment with lysis, similar results were obtained. However, what was unexpected was that Fe@C with similar magnetic characteristics had no visible effect on migration and changes in cell morphology. In this regard, we chose Fe3O4 nanoparticles as the main ones for the study.

### Experiment with changing cell morphology and accelerated migration

A series of in vitro experiments were carried out to demonstrate the possibility of changing cell morphology, orientation and their migration speed in the direction of a constant magnetic field gradient. In a magnetic field, visible aggregates and individual magnetic nanoparticles have a mechanical effect to cytoskeleton through specifically associated with the complex the magnetic nanoparticles (Fe3O4)-dye cy5-ProteinG-antibodies.

The choice of acetylated antibodies to tubulin is determined by the fact that acetylated forms of tubulin are associated with stabilizing modifications of microtubules [10.1073/pnas.1900441116]. Dansylcadaverine, which suppresses vesicular transport in cells [10.1073/pnas.79.7.2291], was used to suppress the endocytosis of nanoparticles. After pretreatment with dansylcadaverine in cells, the nanoparticles stabilized and aggregated to a lesser extent along magnetic field lines. This is important for the specific binding of cytoskeletal fibrils through antibodies present on this complex. Thanks to a strong vertical magnetic field, the complex with magnetic nanoparticles is “dragged” into the cells and helps bypass the endosomal pathway.

It was shown that in a lateral magnetic field, cells with nanoparticles migrate rapidly towards the magnet. Due to the impossibility of automatically photographing cells at certain time intervals, we present the results at the starting and ending points. Accelerated migration in a magnetic field may indicate the presence of a pulling force on the cells towards the magnet. Loading cells with a large number of magnetic nanoparticles without and with antibodies does not affect the degree of deformation in a magnetic field, but accelerates the migration rate.

**Figure 10.**
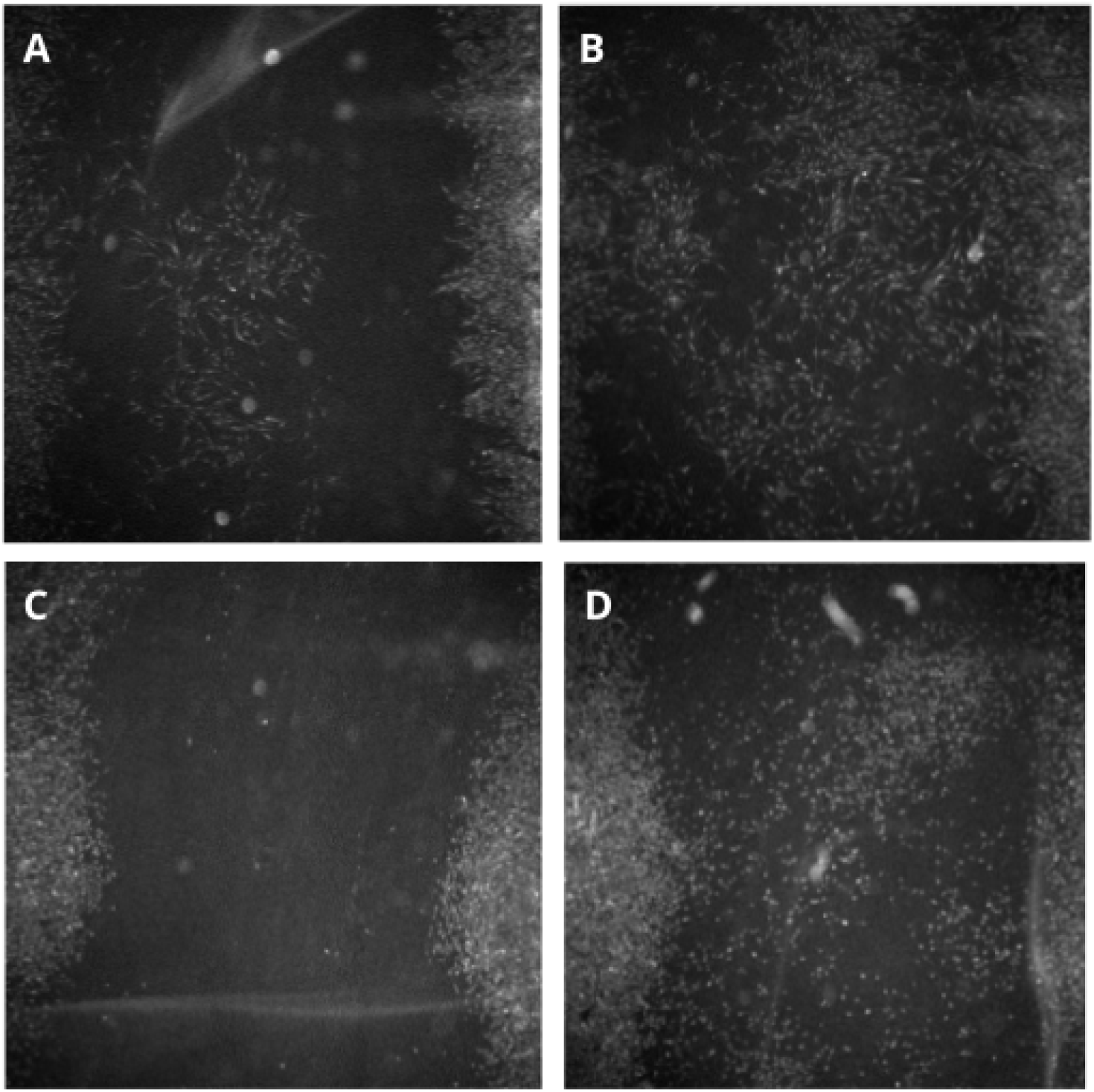
Migration: A – hTert cells, nanoparticle concentration 23.58 μg/ml, B – hTert cells, nanoparticle concentration 117.9 μg/ml, C – MFT-16 cells, nanoparticle concentration 23.58 μg/ml, without magnetic field, D – MFT-16 cells, concentration of nanoparticles 23.58 μg/ml. Cells were incubated with nanoparticles for 24 hours. The magnetic field is directed to the right.

**Figure 11:**
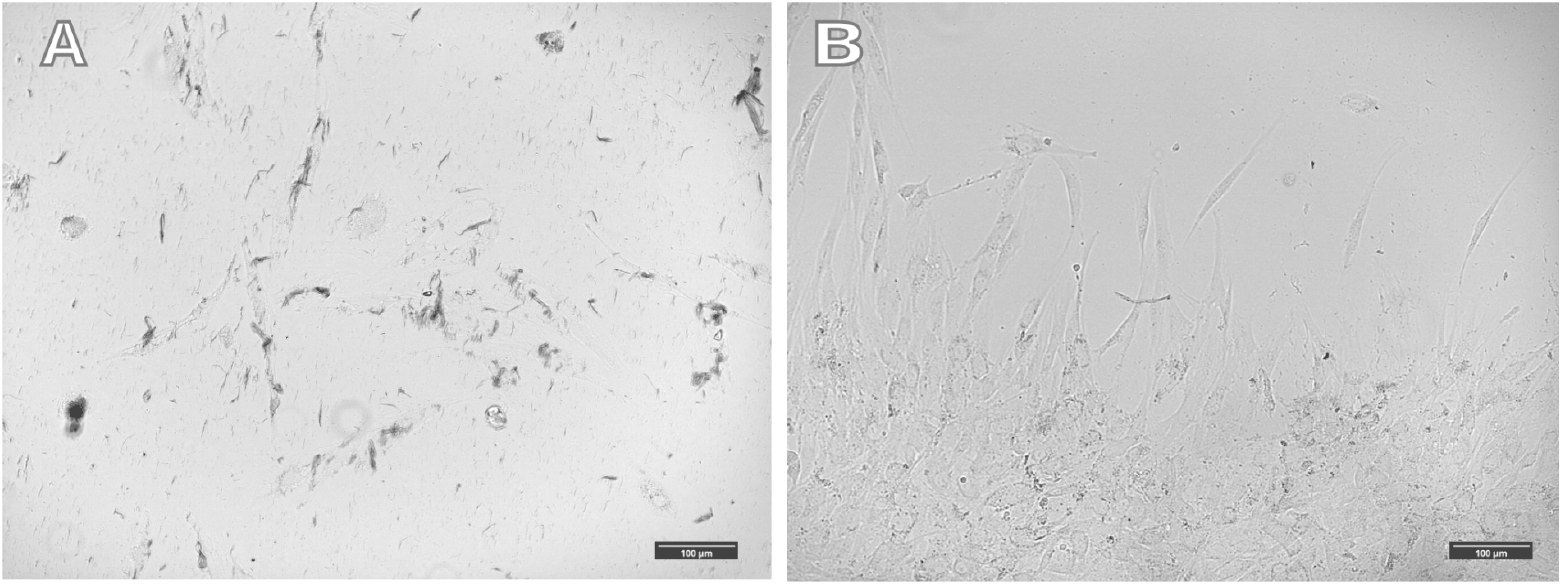
Cells were incubated with different amount of magnetic nanoparticles and then placed in a magnetic system with a horizontal magnetic field for 20 hours. A - concentration of nanoparticles 182.16 μg/ml, B - concentration of nanoparticles 26.2 μg/ml. The magnetic field is directed upwards.

Interestingly, the relationship between the number of nanoparticles and the degree of change in cell morphology has not been reproducibly demonstrated. An increase in the number of nanoparticles in cells does not lead to pronounced changes in cell morphology in a magnetic field. Even filling most of the volume of the cytoplasm with magnetic nanoparticles does not lead to pronounced effects of the magnetic field on cell morphology and the cells are not drawn towards the magnetic field. In such cells, large aggregates of magnetic nanoparticles are observed, aligned along the magnetic field lines. All visible clusters of nanoparticles are their aggregates; single nanoparticles are not visible under an optical microscope.

In a magnetic field, cells with magnetic nanoparticles associated with the cytoskeleton migrate rapidly towards the magnet, but are not drawn towards the magnet and predominantly do not change morphology.

**Figure 12.**
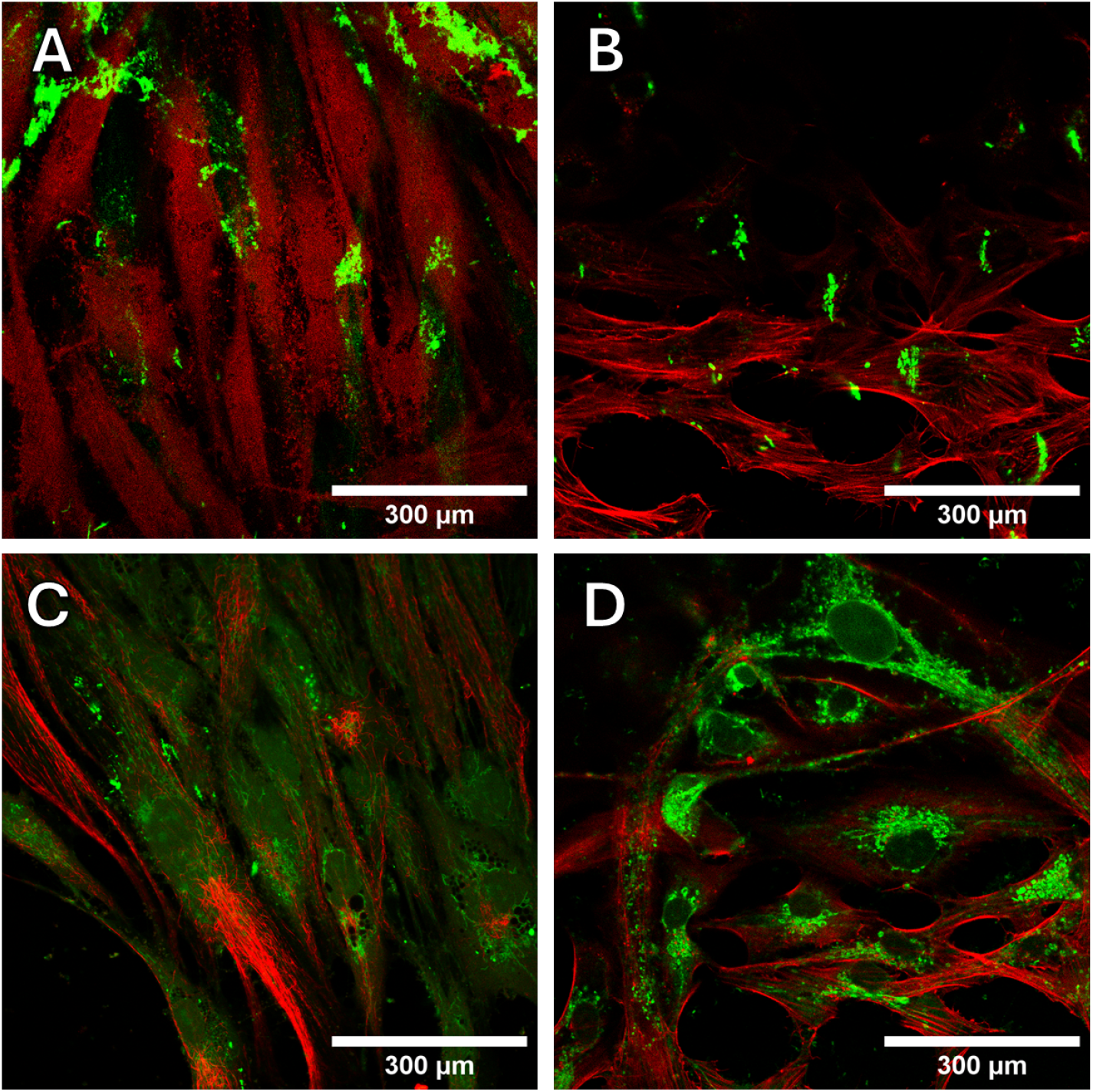
Confocal microscopy of cells incubated on a magnetic system with a horizontal magnetic field. Cells were incubated with magnetic nanoparticles conjugated with different antibodies and Cy5 fluorophore. Immunolabeling with secondary antibodies. A - antibodies to vimentin, B - antibodies to actin, additional staining with phalloidin, C - antibodies to tubulin, D - antibodies to actin, without magnetic field. The magnetic field is directed upwards.

The cells used in the experiment have a good ability to migrate, and to determine the role of the magnetic field and the pulling effect of the complex of magnetic nanoparticles on cells, in a series of experiments we limited the existing ability to migrate. For this purpose, antibodies to Integrin beta-1 were used, which was supposed to ensure “pulling” of cells towards the magnet and limit the ability to migrate. The presence of integrin beta-1 receptor was determined by immunofluorescence. The cells were treated with antibodies to integrin, which slightly limited their migration towards the magnet. However, no pronounced changes in cell morphology were observed (Figure 13).

**Figure 13.**
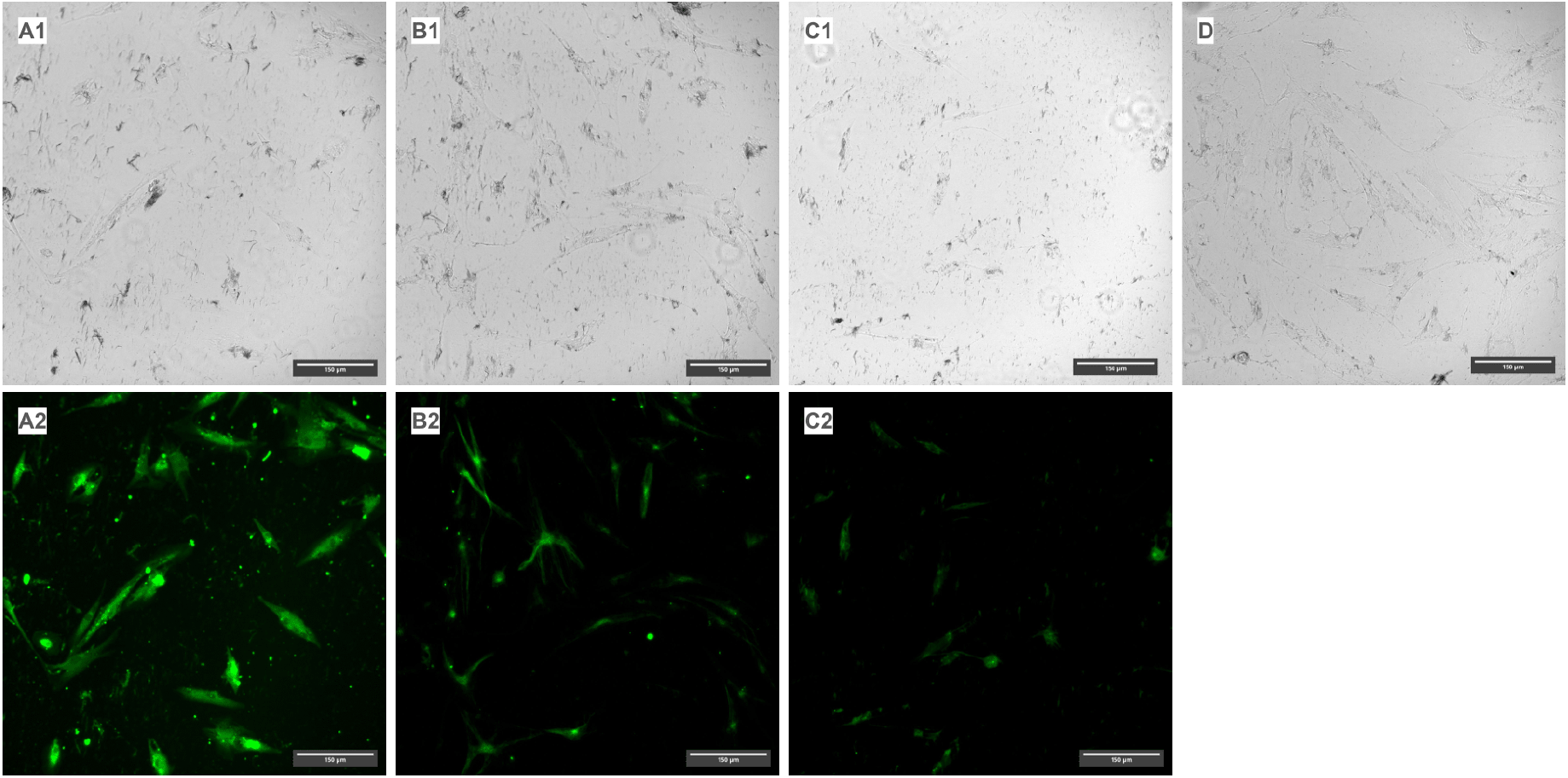
Cells were pretreated with antibodies to β-1 integrin before nanoparticles uptake. Nanoparticles were conjugated to different antibodies. A1-A2: antibodies to beta actin; B1-B2: antibodies to acetylated tubulin; C1-C2 antibodies to vimentin; D: the control sample without antibodies. The magnetic field is directed upwards. Secondary anti-rabbit antibodies labeled with a fluorophore (AF488) were used for visualization.

To reduce the formation of strong contacts with the bottom of the culture vessel and increase the plasticity of cell morphology, an experiment was carried out with reseeding of cells after dragging magnetic nanoparticles into the cells. To capture nanoparticles, 1 ml of nanoparticle solution with a concentration of 182.16 μg/ml was added to the cells. After incubation with nanoparticles for 1 hour, the cells were removed from the plastic and transferred to a new Petri dish and placed on a magnetic system with a horizontal magnetic field for 20 hours. In this way, it is possible to act on the morphology and physiology of cells when their connection with the plastic is not very strong and it is possible to influence the cells. Cells have been shown to adhere to the culture surface but are more aligned with the magnetic field (Figure 14).

**Figure 14.**
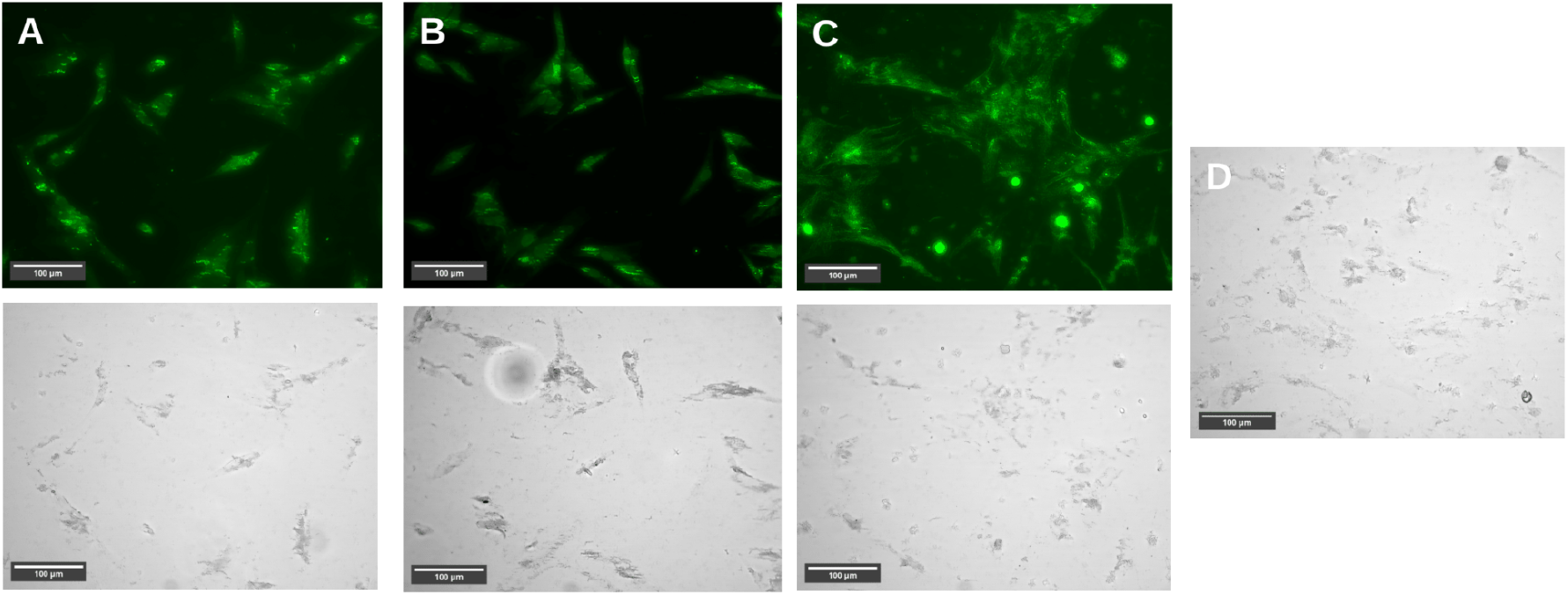
Cells were pretreated with antibodies to β-1 and α-3 integrin, incubated with nanoparticles and reseeding after magnetic nanoparticles uptake. Nanoparticles were conjugated to different antibodies. A - antibodies to actin, B - antibodies to vimentin, C - antibodies to tubulin, D - control without antibodies. The magnetic field is directed to the left. Secondary anti-rabbit antibodies labeled with a fluorophore (AF488) were used for visualization.

## Discussion

There are many works in the literature that show the effect of hyperthermia caused by the influence of an alternating magnetic field on magnetic intracellular nanoparticles. Such work is mainly focused on the treatment of tumor cells through their targeted heating. A study of the influence of a constant magnetic field on biological objects did not show reproducible effects in the absence of magnetic materials in the cells. The question remains open of how a constant magnetic field can affect biological objects with magnetic nanoparticles in cells. Cells in a magnetic field are subject to mechanical forces that pull them towards the magnet due to the presence of magnetic nanoparticles in the cells. Such forces can have two outcomes: pulling cells towards the magnet and accelerated migration.

The first goal of our study was to determine how much antibodies retain their affinity for antigens after cross-linking with nanoparticles. According to paper chromatography, it was shown that antibodies retain the ability to recognize and bind antigens. At the same time, a larger amount of primary antibodies even reduced the intensity of the band, which may be explained by intermolecular cross-links between antibodies on the surface of nanoparticles. At the same time, with direct cross-linking of nanoparticles, dye and primary antibodies, it is not possible to qualitatively visualize the cell cytoskeleton. It is likely that the affinity of the resulting antibodies is not sufficient for labeling individual components of the cell, and such direct cross-linking of nanoparticles and antibodies is not suitable for manipulating cytoskeletal fibrils. In addition, we have shown that indirect conjugation of antibodies with a nanoparticle/dye/protein G or L complex allows efficient binding of their targets in fixed cells and visualization of the cytoskeleton. The good results obtained when using indirect cross-linking of primary antibodies can be explained by the fact that proteins L and G better retain activity after chemical modifications and cross-linking.

As can be seen from the results, the dynamics of the uptake of nanoparticles by cells in the presence and absence of a magnetic field are strikingly different. In the control without a magnetic field, there is a stable, smooth uptake, while in a vertical gradient field, nanoparticles are absorbed by cells during the first 10-15 minutes, after which no noticeable changes in concentration are observed. Subsequently, all cell experiments were carried out after “dragging” nanoparticles into cells using a vertical gradient magnetic field.

An experiment with cell lysis shows the preservation of the affinity of antibodies after cross-linking with nanoparticles and the ability to magnetize cytoskeletal components from cell lysate in a magnetic field. It is important that in such a system there is no viscosity of the cytoplasm and dense intracellular packaging of proteins, which gives greater freedom for manipulating cytoskeletal fibrils. Such a system makes it possible to more freely recognize and form a bond between antibody and antigen; there is no problem of antibody degradation by lysosomal enzymes during endocytosis. We have shown that in the presence of a magnetic field, cytoskeletal components labeled with antibodies, nanoparticles and dye, together with cellular debris, are magnetized towards the magnet. This shows that in the cytoplasm of lysed cells, binding of antibodies cross-linked with nanoparticles to the cytoskeleton and magnetically controlled movement along the field gradient is possible.

The increase in cell migration is possibly due to the pulling effect on the leading edge of the cells and the general mechanical tension towards the magnet. At the same time, adhesion cell cultures cannot exist outside of contact with the bottom of the culture vessel, in suspension. The absence of adhesive contact with the surface is a signal for apoptosis. Contacts with the culture–treated surface of the culture vessel are provided by various integrin protein molecules. Integrins form strong contacts with the substrate and anchor cells [10.1038/s41580-022-00531-5]. It is important that the maximum possible intracellular amount of magnetic nanoparticles is not enough for cells to reproducibly elongate in a magnetic field. When using stronger magnets than presented in our work, the cells are “torn” from the plastic. Statistical analysis of cells shows that cells are oriented along magnetic field lines, and this effect is especially pronounced closer to the magnet. Thus, the cells have only two reactions left - accelerated migration and a change in morphology towards the magnetic field. Due to the fact that animal cells are very plastic and are capable of actively migrating on plastic towards a magnetic field, the degree of their “stretching” may be minimal or absent.

Mechanical tension of the cytoskeletal fibrils deforms the shape of the cell and slightly extends it in the direction of the magnetic field. With a magnetically controlled change in cell morphology, it is important to eliminate the migratory ability of cells, leaving only the ability for mechanical deformation due to the elongation of cells towards the magnetic floor. Depending on which component of the cytoskeleton is associated with magnetic nanoparticles, they cause different cell reactions. The results show that cells with magnetic nanoparticles are able to respond to a magnetic field and slightly change their morphology, stretching towards the magnetic field. These changes are temporary and are observed at the very beginning of exposure of cells to the magnetic field. The reason for these weak changes is the numerous strong contacts with the extracellular matrix and the plastic substrate. It is likely that the mechanical forces generated by magnetic nanoparticles are insufficient for significant and visible deformation in a constant magnetic field.

## Conclusion

Our results open new possibilities for biology and medicine in the field of non-invasive manipulation of cell components and control of cellular processes. An important result of our work was the understanding of how simple tension/stretching of cells on plastic in vitro is limited due to the ability of cells to migrate and strong contacts of integrins with culture plastic. For the first time, we bypassed these limitations and proposed an approach that allows us to influence changes in cell morphology in a magnetic field. Understanding how individual cell components respond to mechanical tension and what signaling and biochemical pathways may be linked will provide new insights into fundamental processes.

## Acknowledgments

The reported study was funded by the Russian Science Foundation Grant #22-74-10041. Confocal microscopic examination was performed using the equipment of the Shared Research Center of Scientific Equipment SRC IIP Ural Branch of RAS. For assistance in the development and construction of the magnetic system and for help with the NMR relaxometry, we express our gratitude to M.A. Uymin, S.V. Zhakov. I.V. Byzov. and Novikov S.I. from the Laboratory of Applied Magnetism of the IMP UrB RAS.

## Notes

### Competing Interest Statement

The authors have declared no competing interest.

## References

1. Lopez, Sara, et al. “Magneto-Mechanical Destruction of Cancer-Associated Fibroblasts Using Ultra-Small Iron Oxide Nanoparticles and Low Frequency Rotating Magnetic Fields.” Nanoscale Advances, vol. 4, no. 2, 2022, pp. 421–36. DOI.org (Crossref), 10.1039/D1NA00474C.

2. Cheng, Dengfeng, et al. “Morphological Effect of Oscillating Magnetic Nanoparticles in Killing Tumor Cells.” Nanoscale Research Letters, vol. 9, no. 1, Dec. 2014, p. 195. DOI.org (Crossref), 10.1186/1556-276X-9-195

3. Wang, Qiwei, et al. “Magnetic Iron Oxide Nanoparticles Accelerate Osteogenic Differentiation of Mesenchymal Stem Cells via Modulation of Long Noncoding RNA INZEB2.” Nano Research, vol. 10, no. 2, Feb. 2017, pp. 626–42. DOI.org (Crossref), 10.1007/s12274-016-1322-4.

4. Labusca, Luminita, et al. “Magnetic Nanoparticles and Magnetic Field Exposure Enhances Chondrogenesis of Human Adipose Derived Mesenchymal Stem Cells But Not of Wharton Jelly Mesenchymal Stem Cells.” Frontiers in Bioengineering and Biotechnology, vol. 9, Oct. 2021, p. 737132. DOI.org (Crossref), 10.3389/fbioe.2021.737132.

5. Labusca, Luminita, et al. “The Effect of Magnetic Field Exposure on Differentiation of Magnetite Nanoparticle-Loaded Adipose-Derived Stem Cells.” Materials Science and Engineering: C, vol. 109, Apr. 2020, p. 110652. DOI.org (Crossref), 10.1016/j.msec.2020.110652.

6. Shen, Yajing, et al. “Cell Mechanosensors and the Possibilities of Using Magnetic Nanoparticles to Study Them and to Modify Cell Fate.” Annals of Biomedical Engineering, vol. 45, no. 10, Oct. 2017, pp. 2475–86. DOI.org (Crossref), 10.1007/s10439-017-1884-7.

7. Cho, Sungwoo, et al. “Tension Exerted on Cells by Magnetic Nanoparticles Regulates Differentiation of Human Mesenchymal Stem Cells.” Biomaterials Advances, vol. 139, Aug. 2022, p. 213028. DOI.org (Crossref), 10.1016/j.bioadv.2022.213028.

8. Gahl, Trevor J., and Anja Kunze. “Force-Mediating Magnetic Nanoparticles to Engineer Neuronal Cell Function.” Frontiers in Neuroscience, vol. 12, May 2018, p. 299. DOI.org (Crossref), 10.3389/fnins.2018.00299.

9. Semeano, Ana T., et al. “Effects of Magnetite Nanoparticles and Static Magnetic Field on Neural Differentiation of Pluripotent Stem Cells.” Stem Cell Reviews and Reports, vol. 18, no. 4, Apr. 2022, pp. 1337–54. DOI.org (Crossref), 10.1007/s12015-022-10332-0.

10. Abdel Fattah, Abdel Rahman, et al. “Targeted Mechanical Stimulation via Magnetic Nanoparticles Guides in Vitro Tissue Development.” Nature Communications, vol. 14, no. 1, Aug. 2023, p. 5281. DOI.org (Crossref), 10.1038/s41467-023-41037-8.

11. Ramazanova, Iliza, et al. “Manipulation of New Fluorescent Magnetic Nanoparticles with an Electromagnetic Needle, Allowed Determining the Viscosity of the Cytoplasm of M-HeLa Cells.” Pharmaceuticals, vol. 16, no. 2, Jan. 2023, p. 200. DOI.org (Crossref), 10.3390/ph16020200.

12. Blümler, Peter, et al. “Contactless Nanoparticle-Based Guiding of Cells by Controllable Magnetic Fields.” Nanotechnology, Science and Applications, vol. Volume 14, Apr. 2021, pp. 91–100. DOI.org (Crossref), 10.2147/NSA.S298003

13. Liu, Jiaojiao, et al. “Manipulation of Cellular Orientation and Migration by Internalized Magnetic Particles.” Materials Chemistry Frontiers, vol. 1, no. 5, 2017, pp. 933–36. DOI.org (Crossref), 10.1039/C6QM00219F.

14. Bongaerts, Maud, et al. “Parallelized Manipulation of Adherent Living Cells by Magnetic Nanoparticles-Mediated Forces.” International Journal of Molecular Sciences, vol. 21, no. 18, Sept. 2020, p. 6560. DOI.org (Crossref), 10.3390/ijms21186560

15. Gao, Jianbo, et al. “Magnetic Field Promotes Migration of Schwann Cells with Chondroitinase ABC (ChABC)-Loaded Superparamagnetic Nanoparticles Across Astrocyte Boundary in Vitro.” International Journal of Nanomedicine, vol. 15, 2020, pp. 315–32. PubMed, 10.2147/IJN.S227328

16. Ho, Vincent H. B., et al. “Generation and Manipulation of Magnetic Multicellular Spheroids.” Biomaterials, vol. 31, no. 11, Apr. 2010, pp. 3095–102. DOI.org (Crossref), 10.1016/j.biomaterials.2009.12.047

17. Perez, Jose E., et al. “Transient Cell Stiffening Triggered by Magnetic Nanoparticle Exposure.” Journal of Nanobiotechnology, vol. 19, no. 1, Dec. 2021, p. 117. DOI.org (Crossref), 10.1186/s12951-021-00790-y

18. Master, Alyssa M., et al. “Remote Actuation of Magnetic Nanoparticles For Cancer Cell Selective Treatment Through Cytoskeletal Disruption.” Scientific Reports, vol. 6, no. 1, Sept. 2016, p. 33560. DOI.org (Crossref), 10.1038/srep33560

19. Královec, Karel, et al. “Disruption of Cell Adhesion and Cytoskeletal Networks by Thiol-Functionalized Silica-Coated Iron Oxide Nanoparticles.” International Journal of Molecular Sciences, vol. 21, no. 24, Dec. 2020, p. 9350. DOI.org (Crossref), 10.3390/ijms21249350

20. Xu, Xueli, et al. “Disturbing Cytoskeleton by Engineered Nanomaterials for Enhanced Cancer Therapeutics.” Bioactive Materials, vol. 29, Nov. 2023, pp. 50–71. DOI.org (Crossref), 10.1016/j.bioactmat.2023.06.016

21. Inaba, Hiroshi, et al. “Magnetic Force-Induced Alignment of Microtubules by Encapsulation of CoPt Nanoparticles Using a Tau-Derived Peptide.” Nano Letters, vol. 20, no. 7, July 2020, pp. 5251–58. DOI.org (Crossref), 10.1021/acs.nanolett.0c01573

22. Erokhin, A. V., et al. “Phenylacetylene Hydrogenation on Fe@C and Ni@C Core–Shell Nanoparticles: About Intrinsic Activity of Graphene-like Carbon Layer in H2 Activation.” Carbon, vol. 74, Aug. 2014, pp. 291–301. DOI.org (Crossref), 10.1016/j.carbon.2014.03.034

23. Postnikov, P. S., et al. “Aryldiazonium Tosylates as New Efficient Agents for Covalent Grafting of Aromatic Groups on Carbon Coatings of Metal Nanoparticles.” Nanotechnologies in Russia, vol. 5, no. 7–8, Aug. 2010, pp. 446–49. DOI.org (Crossref), 10.1134/S1995078010070037

24. Khramtsov, Pavel, et al. “Conjugation of Carbon Coated-Iron Nanoparticles with Biomolecules for NMR-Based Assay.” Colloids and Surfaces B: Biointerfaces, vol. 176, Apr. 2019, pp. 256–64. DOI.org (Crossref), 10.1016/j.colsurfb.2019.01.009

25. Demin, Alexander M., et al. “Silica Coating of Fe3O4 Magnetic Nanoparticles with PMIDA Assistance to Increase the Surface Area and Enhance Peptide Immobilization Efficiency.” Ceramics International, vol. 47, no. 16, Aug. 2021, pp. 23078–87. DOI.org (Crossref), 10.1016/j.ceramint.2021.04.310

26. Demin, Alexander M., et al. “Smart Design of a pH-Responsive System Based on pHLIP-Modified Magnetite Nanoparticles for Tumor MRI.” ACS Applied Materials & Interfaces, vol. 13, no. 31, Aug. 2021, pp. 36800–15. DOI.org (Crossref), 10.1021/acsami.1c07748

27. Marques, A. C., et al. “Functionalizing Nanoparticles with Cancer-Targeting Antibodies: A Comparison of Strategies.” Journal of Controlled Release, vol. 320, Apr. 2020, pp. 180–200. DOI.org (Crossref), 10.1016/j.jconrel.2020.01.035

28. Lunin, Afanasy V., et al. “Synthesis of Highly-Specific Stable Nanocrystalline Goethite-like Hydrous Ferric Oxide Nanoparticles for Biomedical Applications by Simple Precipitation Method.” Journal of Colloid and Interface Science, vol. 541, Apr. 2019, pp. 143–49. DOI.org (Crossref), 10.1016/j.jcis.2019.01.065

29. Girich, Elena V., et al. “Absolute Stereochemistry and Cytotoxic Effects of Vismione E from Marine Sponge-Derived Fungus Aspergillus Sp. 1901NT-1.2.2.” International Journal of Molecular Sciences, vol. 24, no. 9, May 2023, p. 8150. DOI.org (Crossref), 10.3390/ijms24098150

30. Byzov, I. V., et al. “NMR Relaxometry at Quantification of the Captured Magnetic Nanoparticles by Cells.” Physics of Metals and Metallography, vol. 120, no. 13, Dec. 2019, pp. 1341–46. DOI.org (Crossref), 10.1134/S0031918X19130027

31. Minin, A. S., et al. “Application of NMR for Quantification of Magnetic Nanoparticles and Development of Paper-Based Assay.” Journal of Physics: Conference Series, vol. 1389, no. 1, Nov. 2019, p. 012069. DOI.org (Crossref), 10.1088/1742-6596/1389/1/012069

32. Choe, Weonu, et al. “Fc-Binding Ligands of Immunoglobulin G: An Overview of High Affinity Proteins and Peptides.” Materials, vol. 9, no. 12, Dec. 2016, p. 994. DOI.org (Crossref), 10.3390/ma9120994

33. Behzadi, Shahed, et al. “Cellular Uptake of Nanoparticles: Journey inside the Cell.” Chemical Society Reviews, vol. 46, no. 14, 2017, pp. 4218–44. DOI.org (Crossref), 10.1039/C6CS00636A

34. Eshun-Wilson, Lisa, et al. “Effects of α-Tubulin Acetylation on Microtubule Structure and Stability.” Proceedings of the National Academy of Sciences, vol. 116, no. 21, May 2019, pp. 10366–71. DOI.org (Crossref), 10.1073/pnas.1900441116

35. Schlegel, R., et al. “Amantadine and Dansylcadaverine Inhibit Vesicular Stomatitis Virus Uptake and Receptor-Mediated Endocytosis of Alpha 2-Macroglobulin.” Proceedings of the National Academy of Sciences, vol. 79, no. 7, Apr. 1982, pp. 2291–95. DOI.org (Crossref), 10.1073/pnas.79.7.2291

36. Kanchanawong, Pakorn, and David A. Calderwood. “Organization, Dynamics and Mechanoregulation of Integrin-Mediated Cell–ECM Adhesions.” Nature Reviews Molecular Cell Biology, vol. 24, no. 2, Feb. 2023, pp. 142–61. DOI.org (Crossref), 10.1038/s41580-022-00531-5

