## Supplemental materials for "Effect of a constant magnetic field on morphology and motility of cell with cytoskeleton-associated magnetic nanoparticles"

### S1. TEM of magnetic nanoparticles

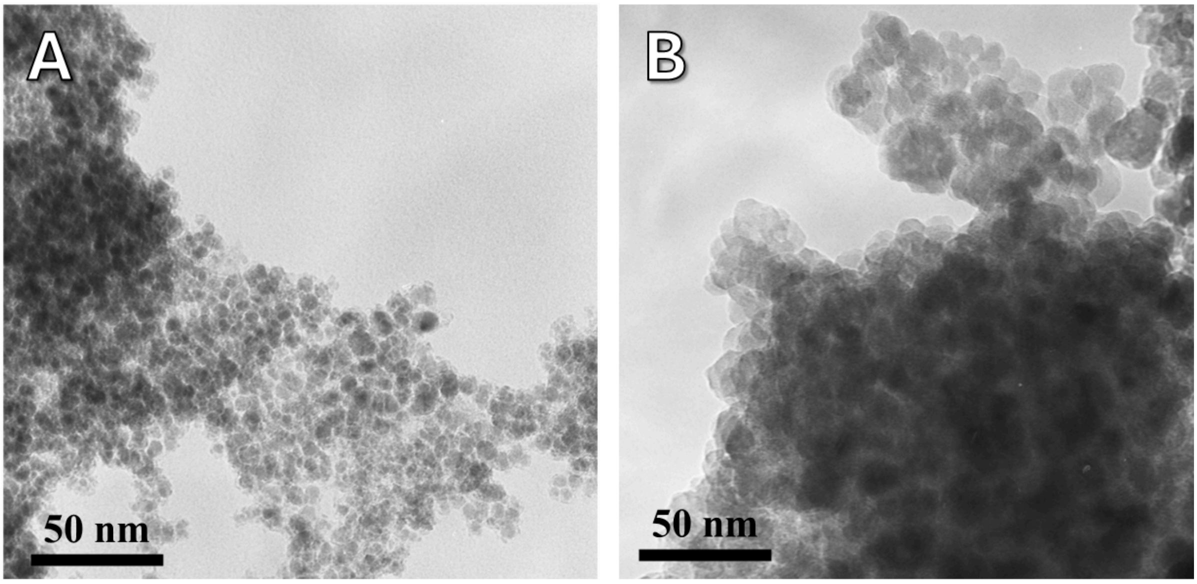


TEM images of a magnetic nanoparticles. A - Fe@C nanoparticles, B - Fe3O4 nanoparticles

### S2. Study of cytoskeleton orientation

The images of cells with actin labelled with Phalloidin-Fluor 633 (Absin, China) taken in different parts of the dish are presented below. The cells were pretreated with antibodies to β-1 integrin for 1 hour (1.5 µl of antibodies were added to 1 ml of culture medium) and then incubated for 1 hour on a vertical magnetic system with nanoparticles cross-linked with antibodies to actin, nanoparticle concentration 182.16 µg/ml, volume of culture medium without FBS 1 ml. After the nanoparticles were captured by the cells, the cell monolayer was washed with a pipette to get rid of the nanoparticles remaining on the cell surface, some of the cells were removed with a scrapper as shown in the figure below, after which the medium was changed to medium with FBS and the dish was incubated on a lateral magnetic system.

The cells were imaged using a confocal microscope and the images were processed in FIJI using the OrientationJ plugin [10.1007/s10237-011-0325-z], which allows the orientation of filaments in the image to be analysed.

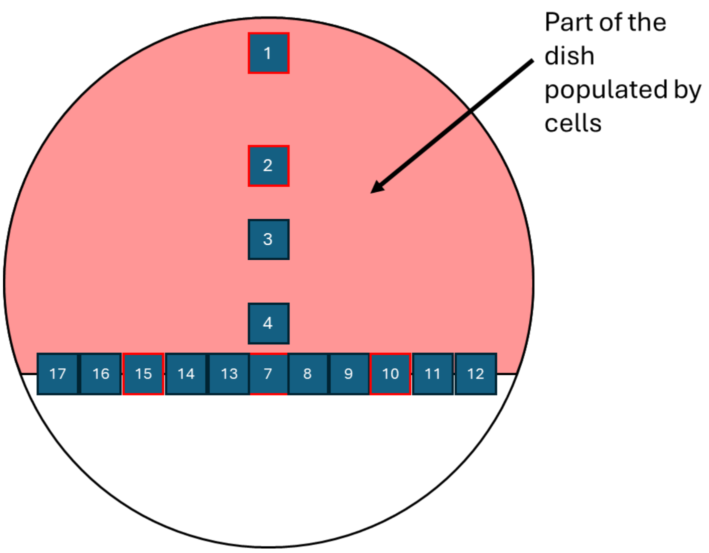


*Map of the working area of the glass-bottomed petri dish in which the cells were imaged. The areas shown in the images below are highlighted in red.*

Below is a typical image, A shows actin filaments, B shows nuclei stained with Hoechst 33258 dye, C shows an image in transmitted light (black spots stand out - clusters of nanoparticles) and D shows an overlay of these images.

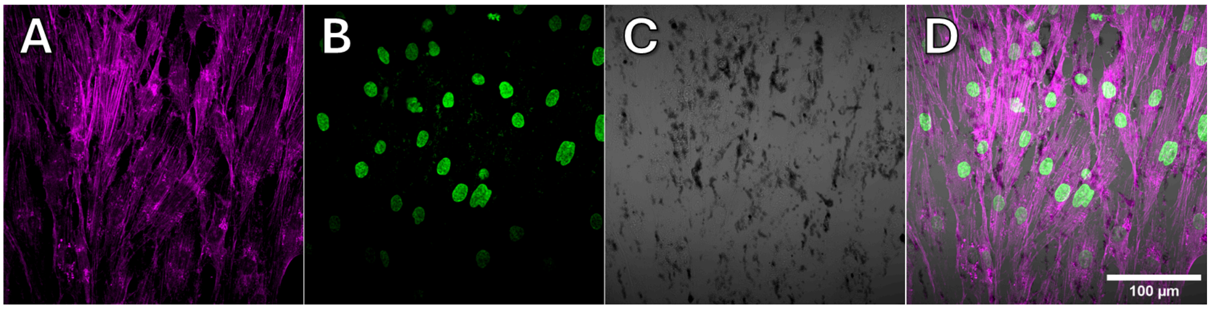


The figures below provide images of actin filaments processed by OrientationJ, coloured in color coordinates where Hue represents filament orientation and saturation represents coherence of nearby filaments, with corresponding filament distribution plots shown next to each other

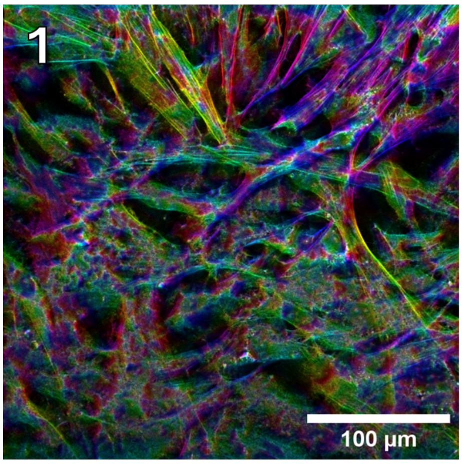
1
100 µm

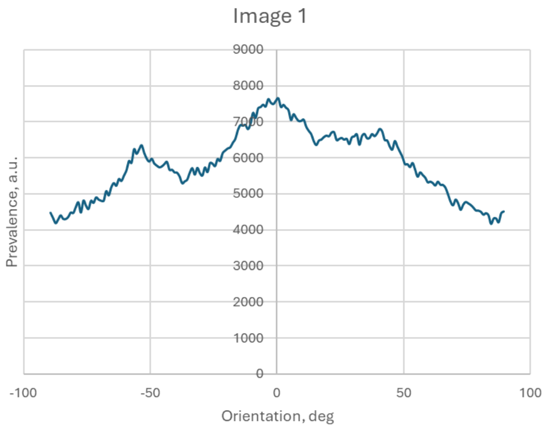
Image 1
9000
8000
7000
6000
5000
4000
3000
2000
1000
0
-100
-50
0
50
100
Prevalence, a.u.
Orientation, deg

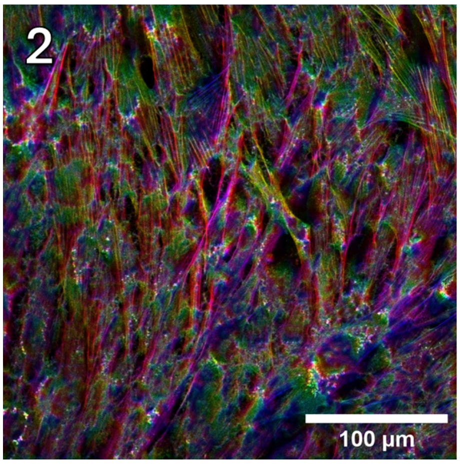
2
100 µm

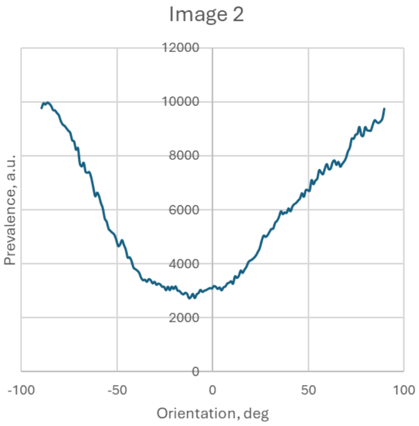
Image 2
12000
10000
8000
6000
4000
2000
0
-100
-50
0
50
100
Prevalence, a.u.
Orientation, deg

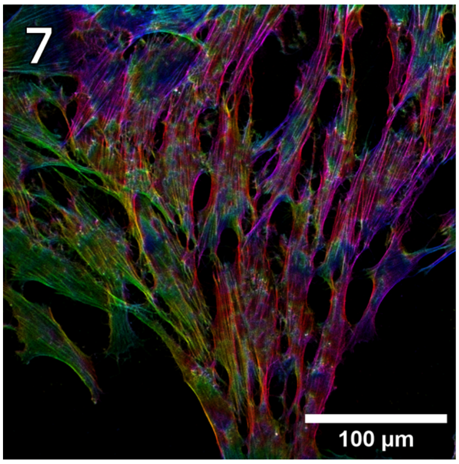
7
100 µm

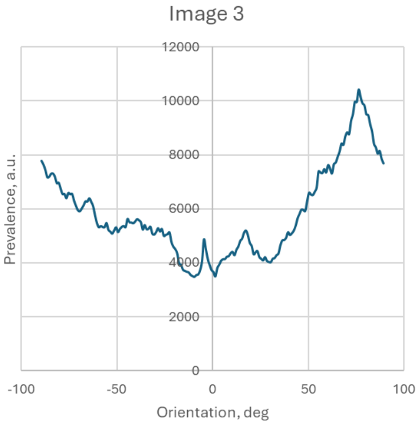
Image 3
12000
10000
8000
6000
4000
2000
0
-100
-50
0
50
100
Prevalence, a.u.
Orientation, deg

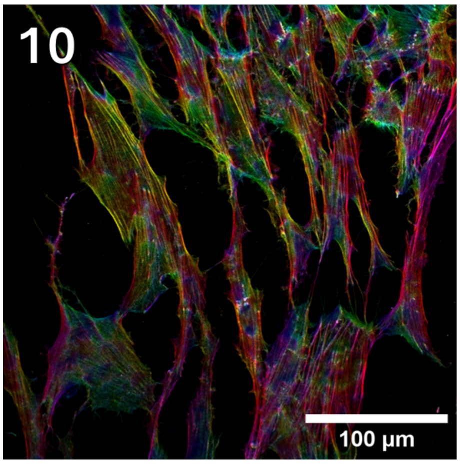


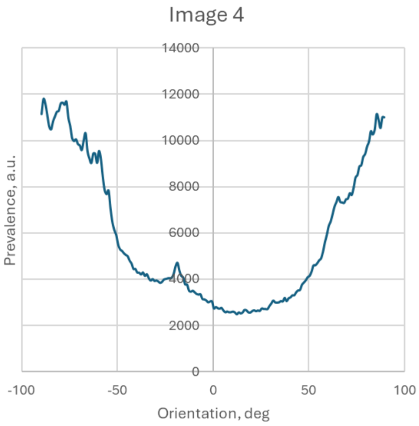


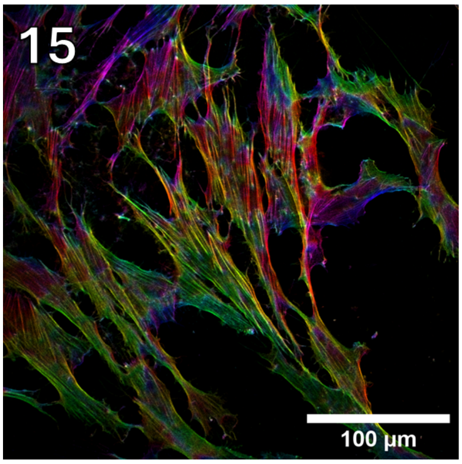


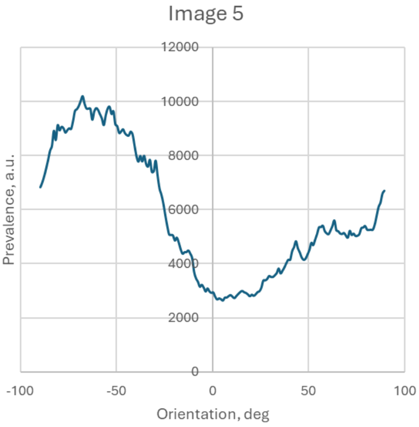


The table below summarises the total figures for filament orientation and coherence in different images

| **Slice** | **Orientation** | **Coherency** |
|---|---|---|
| 1 | 20.295 | 0.051 |
| 2 | 84.71 | 0.16 |
| 7 | 78.138 | 0.286 |
| 10 | 80.092 | 0.371 |
| 15 | 69.091 | 0.315 |

As can be seen from the figures and the table, in the areas where cells grow more freely, as well as closer to the magnetic field where the gradient is somewhat higher, actin filaments begin to predominantly line up parallel to the gradient.

### S3. Antibody affinity testing

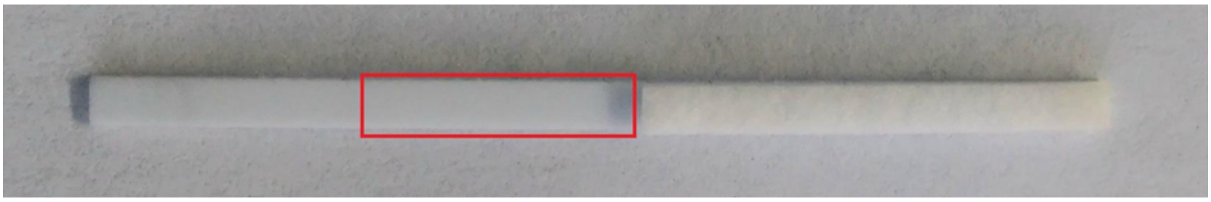

*Figure - Nitrocellulose membrane. The red rectangle indicates boundaries of the BSA drop after application to the centre of the area*

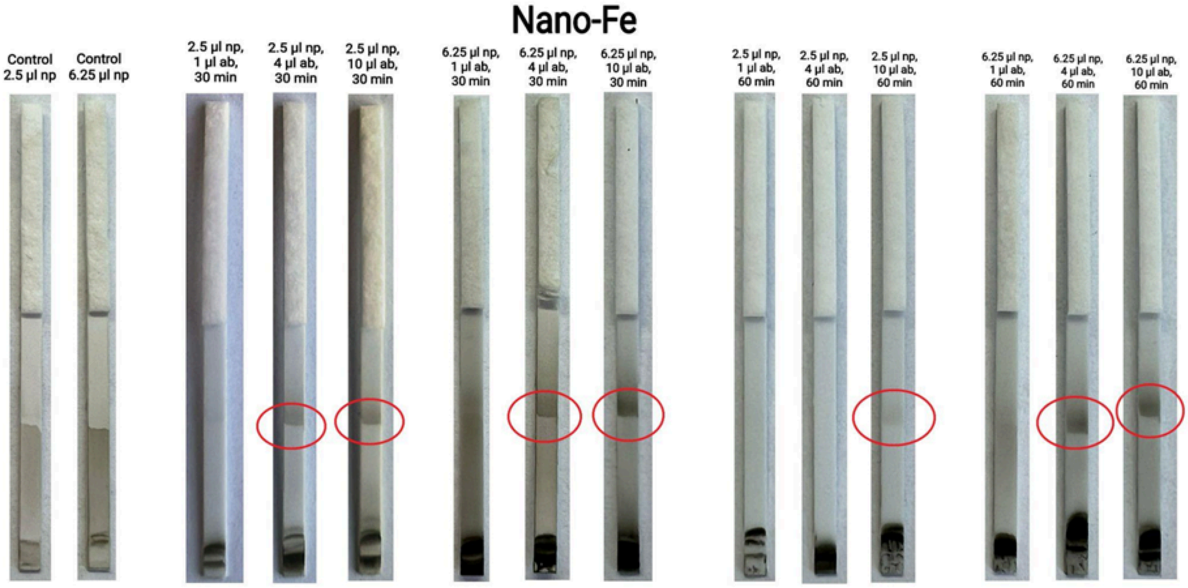


*Figure - Varying crosslinking conditions for n/h Fe (5 mg/ml). The place of contact of nanoparticles with a/g is circled in red.*
